# Changing flows balance nutrient absorption and bacterial growth along the gut

**DOI:** 10.1101/2022.02.16.480685

**Authors:** Agnese Codutti, Jonas Cremer, Karen Alim

## Abstract

Small intestine motility and its ensuing flow of luminal content impact both nutrient absorption and bacterial growth. To explore this interdependence we introduce a biophysical description of intestinal flow and absorption. Rooted in observations of mice we identify the average flow velocity as the key control of absorption efficiency and bacterial growth, independently of the exact contraction pattern. We uncover self-regulation of contraction and flow in response to nutrients and bacterial levels to promote efficient absorption while restraining detrimental bacterial overgrowth.

---

The microbiota of the gut strongly influences intestinal functioning and general health [1–3]. While most bacteria are located in the large intestine [4–9], bacteria are also present in the small intestine (SI) where they exert a strong effect on the host as well. Too high bacterial densities in the SI are particularly problematic as they cause among other symptoms pain, cephalea, chronic fatigue, bloating, and malnutrition [10]. To avoid this *small intestine bacterial overgrowth syndrome* (SIBO) luminal flow and the active transport via gut muscle contractions is essential [10]. Gut motility [11–14] further affects nutrient absorption, while motility patterns vary, with peristalsis prevalent during starvation [11–19] and the ‘checkerboard-like’ segmentation pattern during digestion after food intake [11–14]. From a physics perspective gut motility drives fluid flows [15, 16] and thus impacts the dispersion and transport of solutes [20–23]. Gut motility may therefore well control nutrient absorption and bacterial densities [17] with all processes being highly intertwined. Bacteria for example influence nutrient levels as they compete with the gut for their absorption, and nutrient availability as well as bacterial densities feed back onto gut motility [11, 13, 14, 24, 25] (see Fig. 1A). While motility driven flow [26–40], peristalsis-induced nutrient absorption, [41–43] and bacterial growth [17] have been independently investigated, the complex dynamics arising due to different motility patterns and feedback from bacteria or nutrient density remains unknown.

**FIG. 1.**
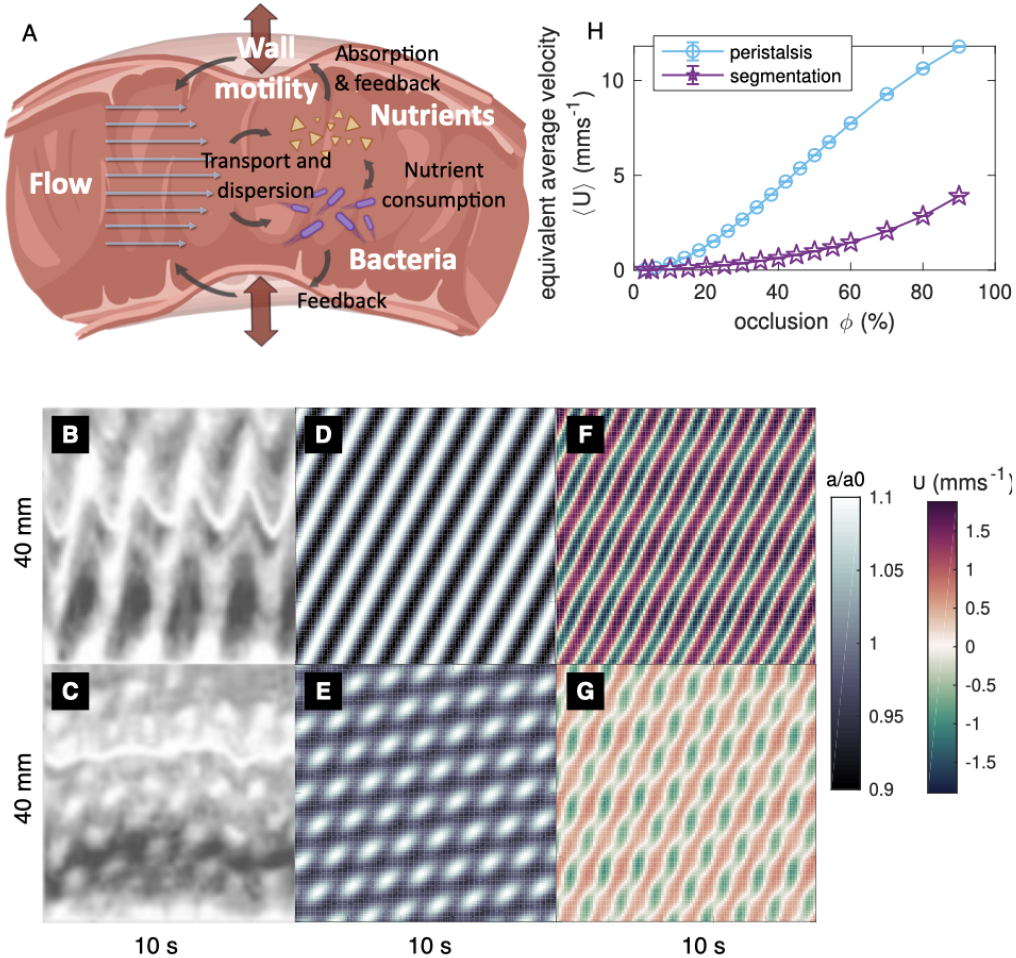
Gut motility determines flows. (A) The gut is a muscular tube, whose motility patterns induce flows that affect the abundance of nutrients and bacteria. Abundances, in turn, feed back on motility. (B, C) *In vitro* spatio-temporal map of the contraction amplitude observed for the small intestine of mice, during peristalsis and segmentation [14], respectively. Data from Huizinga *et al*. [14] with permission. (D, E) Simulated contraction amplitudes *a*(*t, z*)/*a*_0_ with 10% occlusion and (F, G) the emerging flow *U* for peristalsis and segmentation. (H) Equivalent average flow velocity 〈*U*〉 as function of maximal tube occlusion *ϕ* for peristalsis (blue) and segmentation (purple). (A) courtesy of Sara Gabrielli.

Here, we investigate the interdependence of flow, nutrient absorption, and bacterial growth for a diverse set of motility patterns. To gain analytical insights we extend the Taylor dispersion approach [20–23, 44, 45] and setup a model which accounts for spatio-temporally contracting walls, the distribution of nutrients dispersed by the flow and diminished by absorption at the gut wall, and the bacterial growth. We explore the experimentally well-studied mouse gut as a reference scenario and identify the average flow-velocity as the key driver of absorption and bacterial growth dynamics independent of the underlying motility pattern causing flow. In fact, we show that physiological feedback precisely controls flow velocity to balance nutrient absorption and bacterial growth.

To account for the variety of contractility patterns changing over time *t* and along the intestine’s longitudinal direction *z* we describe the variation of the gut radius *a*(*z, t*) around a rest radius *a*_0_ as a superposition of two sine waves with high *H* and low *L* frequencies [14]

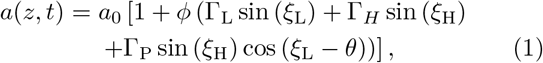

Here, *ϕ* denotes the maximal occlusion, *ξ* ≔ Ω*t* − *Kz*, Ω and *K* are temporal and spatial frequencies, with Ω_H_ > Ω_L_ and *K*_H_ > *K*_L_, *θ* is the phase shift between the high and low frequency wave, and Γ are generic coefficients normalized such that the factor multiplied with *ϕ*, i.e. the overall occlusion depth, is at maximum 1. Two prominent contractility patterns observed in mice [12] are represented by this function, i.e. peristalsis for Γ_P_ = Γ_L_ = 0, Γ_H_ = 1 (Fig. 1(B,D)); and segmentation for all coefficients non-zero Γ_P_ = 0.48, Γ_L_ = 1, Γ_H_ = 0.78 [14] (Fig. 1(C,E); see also I [46]). For the long slender geometry of the small intestine the contractility driven flow of velocity *U* is described by Stokes flow following directly from the spatio-temporal contractions of the tube, the applied pressure drop Δ*p* along the tube of length *L* and the fluid viscosity *μ* [16] (see II [46]). To describe nutrient *N* and bacteria concentration *B* we assume that flow in the gut is quasi laminar, i.e. *a*_0_ << *K*/(2*π*), that concentration gradients across the tube’s cross-section average out quickly by diffusion with diffusion coefficient *κ*, i.e. 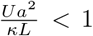, and that nutrient absorption is small, i.e. *γa*/*k* < 1 with *γ* being the absorption strength. These conditions are approximately met for experimental parameters derived from the mouse model [14, 17, 47–50], under the assumption of small occlusion and water viscosity (see I, III [46]). Therefore, we derive the spatio-temporal dynamics for the cross-sectionally averaged concentrations of nutrients and bacteria within the framework of Taylor dispersion employing the invariant manifold method [20–23, 44, 45] while accounting for an absorbing tube wall undergoing spatio-temporal contractions (derivation in IV, numerical details in V [46]). Using Monod kinetics to describe the bacterial growth [17, 51, 52], the dynamics of the nutrient concentration *N* is

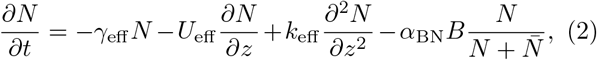

where 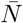 denotes the nutrient concentration below which growth is hindered [17, 51, 52]. The effective components are:

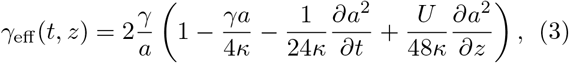

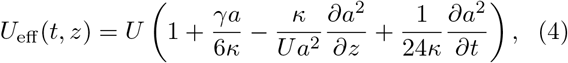

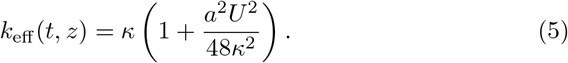

The corresponding equation for bacteria *B* is

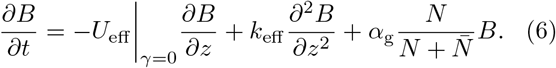

To assess the effect of gut contractility on ensuing flow, we consider the two prominent contractility patterns introduced before, peristalsis (see Fig. 1B, D) and segmentation (Fig. 1C, E). At equal tube occlusion, peristalsis produces stronger and more persistent longitudinal flows (both anterograde and retrograde, Fig. 1F) than segmentation (Fig. 1G). The equivalent average flow velocity over a period of contraction 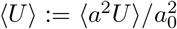 (*i.e*. the flow velocity that an equivalent straight tube with the same volumetric flow would have II [46]), increases with tube occlusion and is also much stronger for peristalsis than for segmentation (Fig. 1H). Notably, slowing down peristalsis to achieve a lower 〈*U*〉 is not equivalent to employing segmentation, since in the latter case longitudinal flows *U* are strong and occlusion *ϕ* is high, which is implicated in enhancing mixing [34, 35, 40, 53–56]. In conclusion, the gut has different controls of flow velocity by either adapting the muscles strength that is coordinating tube occlusion or by retaining the same occlusion but modifying the spatio-temporal pattern of contractions, i.e. changing the wave superposition in Eq. 1.

To determine how different flow patterns impact nutrient dispersion and absorption, we follow the spread of a finite amount of nutrients normally distributed at time zero with free outflow and inflow, *∂N*/*∂z*|_boundary_ = 0, first in the absence of bacteria. In agreement with experimental observations [57], the nutrient dispersion is directly modulated by the contraction patterns as illustrated by the outflow behavior shown in Fig. 2A. Yet, we observe that the residence time *t*_res_ ≔ *∫* d*t* (*t* · *J_N_*|_*L*_) / *∫* d*tJ_N_*|_*L*_, with 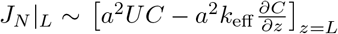 the approximated flux at the outlet [23] (see VI [46]), is independent of the local variations in the flow field incorporated by the different patterns, but only depends on the equivalent average flow velocity: *τ*_res_ = *L*/(2〈*U*〉), see Fig. 2B. The same applies for the nutrient absorption rate Φ_N_ ≔ − *∫_S_*(*k*∇*N*)_⊥_d*S* (mol s^−1^) across the entire tube’s surface *S* (VII [46]), whose decay rate *τ*_abs_, *i.e*. the absorption time, is derived analytically and is in good approximation 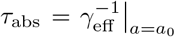 for all motility patterns (see VIII, IX [46]). Therefore, the efficiency, defined as total amount of absorbed molecules until the tube empties normalized by the initial amount of molecules *∫*_5*τ*_res__ Φ_N_d*t*/*N*_initial_, is also pattern independent, see Fig. 2C.

**FIG. 2.**
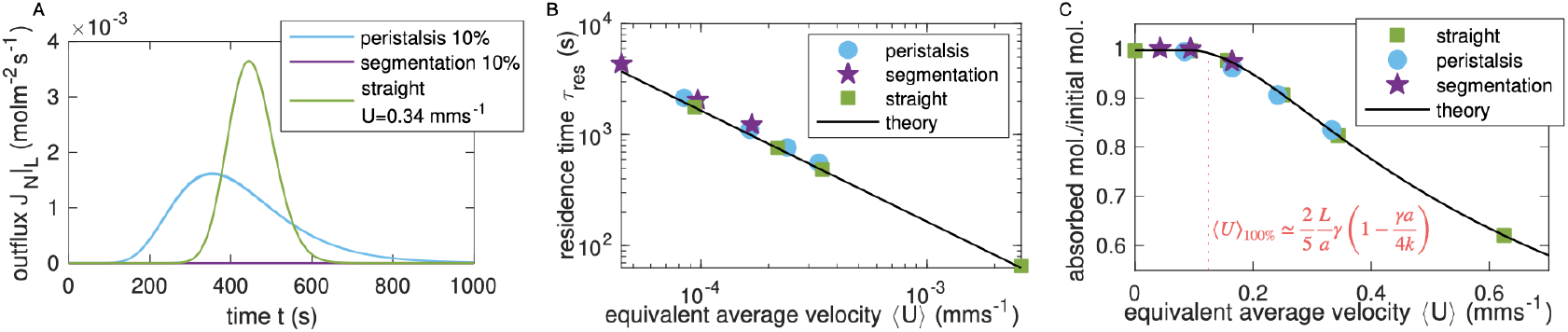
Flow velocity governs residence times and nutrient absorption. A) Average out-flux of nutrients for peristalsis 10% occlusion (light blue), segmentation 10% occlusion (purple), and a straight tube with the 10%-peristalsis-equivalent average flow velocity 〈*U*〉 (green). B) Residence times *τ*_res_ as function of equivalent average flow velocity 〈*U*〉 for peristalsis (light blue), segmentation (purple), and straight tube (green). C) Total absorbed molecules during emptying time normalized by the initial molecules in the tube as function of the equivalent average flow velocity 〈*U*〉 for peristalsis (light blue), segmentation (purple), a straight tube (green), and theoretical prediction for a straight tube (black). The vertical line is the theoretical velocity 〈*U*〉_100%_ above which there is no complete absorption.

Given this result we deduce that, for small velocities 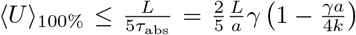 that allow for the residence time to be longer than the absorption time *τ*_res_ > *τ*_abs_, 100% nutrient absorption efficiency can be reached, independently of the flow-generating contractility pattern (red vertical line in Fig. 2C). Thus, when considering only the gut’s role to absorbe nutrients, low flow velocities below 〈*U*〉_100%_ seem ideal [11, 27]. Yet, we have so far neglected the impact of flow on bacterial concentration.

Modeling the stomach as an upheld reservoir of a fixed concentration of bacteria and nutrients *N*|_0_= *N*_0_, *B*|_0_ = *B*_0_ and allowing free outflow *∂N*/*∂z*|_L_ = 0 [17], we analytically solve for both nutrient and bacteria dynamics at steady state 〈*∂N*/*∂t*〉 = 0, 〈*∂B*/*∂t*〉 = 0 for a straight tube. Employing that wall absorption dominates over bacterial consumption terms in Eq. 2 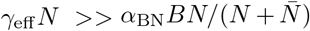 and neglecting the diffusivity term (as confirmed by simulations in X [46]), we obtain

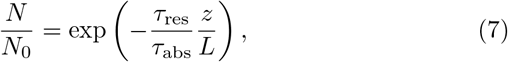

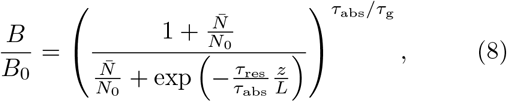

where we further approximated *U*_eff_ ~ *U* = *L*/*τ*_res_ given the small absorption strength *γa*/*k* < 1, and where we defined the growth time as 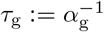. This analytical result, in qualitative agreement with previous simulations [17], clearly states that nutrient concentration is regulated by the competition between advection and absorption timescales. Bacterial concentration is additionally controlled by the competition between nutrient absorption *τ*_abs_ and bacterial growth itself set by the timescale *τ*_g_, with high numbers of bacteria arising for large *τ*_abs_/*τ*_g_, *i.e*. when bacteria multiply much faster than the depletion of nutrients via absorption.

Which of these timescales *τ*_g_, *τ*_abs_ and *τ*_res_ does the gut regulate to improve absorption and limit bacterial growth? For a finite amount of nutrients inside the gut we beforehand found that the residence time, which is governed by the equivalent average flow, is the most important timescale determining absorption (Fig. 2C). Indeed, this is also true here, independent of the motility pattern, for the steady state with an upheld concentration of both nutrients and bacteria as confirmed by simulations. The equivalent flow velocity is regulating the absorption rate Φ_N_ (Fig. 3A; rate normalized by the straight-tube infinite-velocity limit Φ_Uinf_ = 2*πkLC*_0_*γ*(1 − *γ*/4)(1+*γ*/6)(1+*γ*/12)^−1^); see also X [46]). In apparent contrast to the case of a finite amount of nutrients, higher flows correspond to better absorption. However, this is due to the higher inflow of nutrients into the gut when flow increases. Indeed, the efficiency given as the absorption rate normalized by the influx 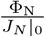 confirms our previous result that higher flows have a much lower efficiency, reaching 100% only for low velocities, also in agreement with Ref. [27]. Flow is also regulating overall bacterial abundance 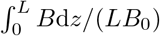 (Fig. 3B and X [46]), hindering it at high velocities and favoring it at small velocities. We further confirm that nutrient absorption and bacterial growth are largely pattern-independent (Fig. 3A and B), with good agreement between simulations and predictions by the straight-tube theory, and that the dynamics of the system is again dominated by the equivalent average flow as key parameter. Another regulatory parameter is the absorption strength *γ*, with higher *γ* promoting higher absorption efficiency (Fig. 3A) and less bacterial growth (Fig. 3B). Notably, the physiological absorption strength *γ* = 10^−6^ ms^−1^ [17] retains an efficiency of at least 40% even for the unfavorable case of fast flows, while it strongly limits bacterial growth, never surpassing 270% even when flow is slow.

**FIG. 3.**
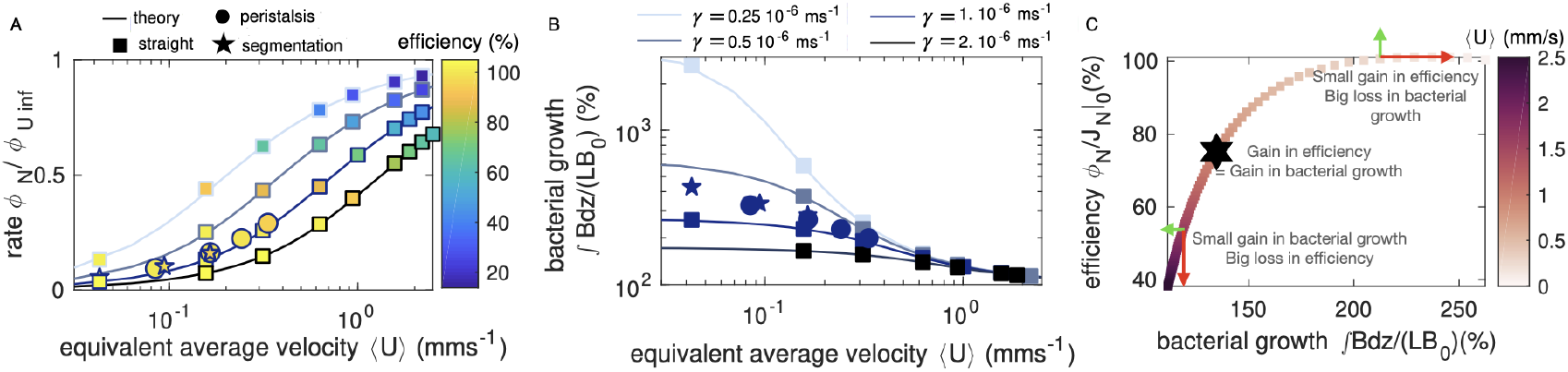
Nutrients absorption and bacterial growth need to be balanced. Comparison between straight tube theory (lines) and simulated (squares for straight tube, circle for peristalsis, and pentagram for segmentation) of the A) normalized absorption rate Φ_N_/Φ_U inf_ and B) bacterial growth 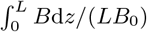 as function of the equivalent average flow velocity 〈*U*〉, at steady state with upheld concentration at the inlet. Different absorption strength *γ* are given (from light blue to dark blue, respectively 2 · 10^−6^ ms^−1^, physiological parameter 1 · 10^−6^ ms^−1^, 0.5 · 10^−6^ ms^−1^, 0.25 · 10^−6^ ms^−1^). In A), data points are color-coded by the steady state efficiency Φ_N_/(*J_N_*|_0_). C) Theoretical steady state efficiency vs. bacterial growth for a straight tube with upheld concentration, color coded by the equivalent average flow velocity 〈*U*〉. The hexagram is the optimal point with flow 〈*U*〉 = 0.88 mm s^−1^, simultaneously optimizing the efficiency (74%) and the bacterial growth (134%).

Taken together, the small intestine of mice appears to operate in a parameter range where changing the motility pattern, and thus the flow velocity, is an efficient mechanism to promote nutrient absorption and limit bacterial growth. Yet, does an optimal flow velocity exist? Plotting the theoretical steady state efficiency as a function of bacterial growth for a straight tube with upheld concentration Fig. 3C, we define the optimum as the point where the curve derivative *δ*efficiency/*δB* is equal to 1 (at 〈*U*〉 = 0.88 mm s^−1^). Moving from that point by increasing flow slightly favors bacterial reduction but significantly worsens nutrient absorption, with the opposite when flow decreases. Thus, while this optimal velocity does not fully optimize both aims, it might provide an acceptable value for both of them simultaneously (74% absorption efficiency and 134% bacterial growth). These results also suggest that a better strategy might be to alternate between two phases of flow, with slow flow during segmentation to fully optimize absorption, and fast flows during peristalsis to down-regulate bacteria. Experimental observations [11, 12, 27, 58] support such alternations in coordination with meal-intake and fasting.

It remains an open question whether the switch between segmentation (of duration *T*_abs. phase_) and peristalsis (duration *T*_clean phase_) is rather driven by nutrient availability [11, 13, 14, 24] or bacterial abundance [25]. From the perspective of maximizing absorption after a meal, flow needs to be slow enough 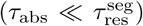 for a long enough duration (*τ*_abs_ ≪ *T*_abs. phase_) to maintain nutrients in the gut and allow absorption to take place. To subsequently clean the gut from bacteria, flow needs to be fast enough such that the residence time is shorter than the bacterial growth timescale 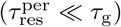 and last long enough to flush out most bacteria 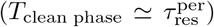. In principle, a very long absorption phase can clean out bacteria, since eventually nutrients are so scarce that bacterial growth is inhibited and bacteria are flushed out, see Fig. 4. This is, however, a very risky strategy as an early arrival of a new meal before the completion of such a slow washout may result in bacterial overgrowth. Instead of being indefinite, the absorption phase is, thus, only maintained temporarily and the flow-pattern switch is coordinated depending on the state of the system. In particular, if bacterial growth is very slow compared to the absorption and residence timescales (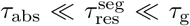 or 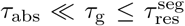), the absorption phase maximizes the efficiency while keeping at bay bacterial growth if it lasts 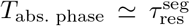. In this scenario a feedback control in which the depletion of nutrients triggers peristaltic cleaning appears to be sufficient to quickly reduce bacteria while ensuring high efficiency, see Fig. 4. If instead bacterial growth is very quick 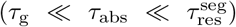, overgrowth is eminent if 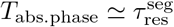. Here, a feedback control in which high bacterial densities trigger peristalsis and limit absorption phase below a duration 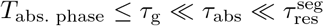 is required to limit bacteria growth. This comes at the cost of reduced efficiency, which can be counteracted by maximizing 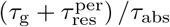 (XI [46]). For the healthy mouse gut, *τ*_abs_ < *τ*_g_ holds (III [46]) and a feedback control via nutrients appears to be efficient. However, disease and other disruptions may affect this parameter balance and require a feedback control via bacteria - at least as a back-up option.

**FIG. 4.**
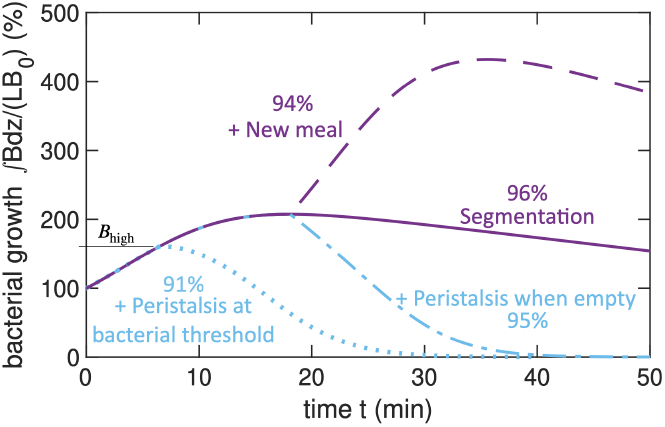
Alternating patterns improve efficiency and bacterial regulation. Absorption phase (segmentation 10%, initially uniformly nutrient and bacteria-filled tube, no inflow, free outflow, purple filled line, see XI [46]) reaches 96% of efficiency. Adding a meal (at 18 min) triggers strong bacterial growth (purple dashed line). Switching to fast flows (peristalsis 10%) when the tube is empty (94% of absorbed molecules, peak of bacterial population) quickly decreases the bacterial number (blue mixed line). Starting peristalsis above a threshold of bacterial growth *B*_high_ = 260% implies higher loss of efficiency at 91% (blue dotted line).

In conclusion, we introduced a coarse-grained analytical description to investigate the complex interdependence of motility, flow, nutrient absorption, and bacterial growth along the small intestine. Our analytical insight allowed us to gain a mechanistic understanding of how flow-control by feedback mechanisms promotes nutrient absorption and prevents bacterial growth. The specific mapping to experimental measures of simple timescales provides a direct interface to experiments and together with the theory promotes an integrative understanding of the intestine and its physiological processes. Society.

## ACKNOWLEDGMENTS

We thank Sophie Marbach for discussions on Taylor dispersion. The work was supported by the Max Planck

## Notes

### Competing Interest Statement

The authors have declared no competing interest.

