## Supplementary Information for "Changing flows balance nutrient absorption and bacterial growth along the gut"

### Supplementary materials for Changing flows balance nutrient absorption and bacterial growth along the gut

(Dated: February 15, 2022)

In this file, we provide the additional derivations and data to support our main manuscript.

#### I. ANALYTICAL PATTERN

The segmentation is inspired by the work of Huizinga [1]. In their work, they show that segmentation is a peristaltic wave with high frequency (in their paper called MP wave) modulated by another wave at low frequency (DMP wave). For segmentation alone, only the high-frequency wave remains. Here, we simplify and generalize their description of segmentation, to have an effective model. Segmentation is thought of as a modulation of the radius $a$ in time and space:

$$a_{\mathrm{segmentation}}(t,z) = a_0\left(1+\phi\,\mathrm{signal}(t,z)/\mathrm{renorm.}\right). \tag{1}$$

The signal is the modulation which in Huizinga 2015 is called $f_{\mathrm{sum}}$, so we can write, from their Equation 1

$$\mathrm{signal} = f_{\mathrm{L}} + f_{\mathrm{H}} \tag{2}$$

where we used the notation L for the DMP signal and H for the MP signal. From their Equation 2, we get for the low frequency signal:

$$f_{\mathrm{L}} = \frac{A_{\mathrm{L}}}{2}\sin\left(\omega_{\mathrm{Lt-K_Lz}}\right) := \Gamma_{\mathrm{L}}\sin(\xi_{\mathrm{L}}) \tag{3}$$

where we renamed for simplicity the phase $\xi_{\mathrm{L}} := \omega_{\mathrm{Lt-K_Lz}}$ and the amplitude $\Gamma_{\mathrm{L}} := \frac{A_{\mathrm{L}}}{2}$. They observe that the high frequency amplitude therefore is modulated by the low frequency phase. From their Equation 3, putting the baseline to 0 as they do throughout the paper, therefore they write the high frequency signal as

$$f_{\mathrm{H}} = \frac{A_{\mathrm{H}}(\tilde{\xi}_{\mathrm{L}})}{2}\sin\left(\omega_{\mathrm{Ht-K_Hz}}\right) := \frac{A_{\mathrm{H}}(\tilde{\xi}_{\mathrm{L}})}{2}\sin(\xi_{\mathrm{H}}) \tag{4}$$

where $\tilde{\xi}_{\mathrm{L}}$ is the bounded phase of the low frequency signal, which they define as a bounded value between 0 and $2\pi$ as:

$$\tilde{\xi}_{\mathrm{bounded}} = \pi - 2\tan^{-1}\left[\cot\left[\frac{1}{2}\left(\omega t - kz + \frac{\pi}{2}\right)\right]\right]. \tag{5}$$

Since for our purposes we do not need a bounded value between 0 and $2\pi$ (we will use sine functions which are bounded while they use exponential functions which need to be bounded by hand), we use the following equivalent function as the phase of the low-frequency signal:

$$\tilde{\xi}_{\mathrm{bounded}} \to \tilde{\xi}_{\mathrm{L}} = \omega_{\mathrm{L}}t - K_{\mathrm{L}} + \frac{\pi}{2} = \xi_{\mathrm{L}} + \frac{\pi}{2} \tag{6}$$

which is equivalent to their definition.

In their Equation 5, they propose the amplitude of the high-frequency signal to depend on the low-frequency signal through a sum of Gaussians. We aim to simplify, so we use the fit they do on their experimental data, where it can be seen that

*

$$2A_{\mathrm{H}}(\tilde{\xi_{\mathrm{L}}}) = B + A\sin\left(\tilde{\xi}_{\mathrm{L}} - \theta\right) \tag{7}$$

which substituting our equivalent phase definition becomes

$$2A_{\mathrm{H}}(\tilde{\xi_{\mathrm{L}}}) = B + A\cos\left(\xi_{\mathrm{L}} - \theta\right). \tag{8}$$

The values A and B can be found fitting Figure 2E from Huizinga [1], as rough estimate A=1.9 mV and B=3.1 mV.

To get a segmentation pattern comparable to the figure of figure 4E in Huizinga [1], where the bright spots are well defined, we set the shift $\theta$ to $0.3 \cdot 2\pi$ rad.

Using Supplementary Eq. 2, 3, 4, and 8 we obtain:

$$\mathrm{signal}(t,z) = f_{\mathrm{L}} + f_{\mathrm{H}} = \Gamma_{\mathrm{L}}\sin(\xi_{\mathrm{L}}) + \frac{A_{\mathrm{H}}(\tilde{\xi_{\mathrm{L}}})}{2}\sin(\xi_{\mathrm{H}}) = \Gamma_{\mathrm{L}}\sin(\xi_{\mathrm{L}}) + \frac{B + A\cos\left(\xi_{\mathrm{L}} - \theta\right)}{4}\sin(\xi_{\mathrm{H}}) \tag{9}$$

Thus, renaming $B/4 := \Gamma_{\mathrm{H}}$ and $A/4 := \Gamma_{\mathrm{p}}$, a general expression for the segmentation signal is found:

$$\mathrm{signal}(t,z) = \Gamma_{\mathrm{H}}\sin\left(\Omega_{\mathrm{H}}t - K_{\mathrm{H}}z\right) + \Gamma_{\mathrm{L}}\sin\left(\Omega_{\mathrm{L}}t - K_{\mathrm{L}}z\right) + \Gamma_{\mathrm{P}}\sin\left(\Omega_{\mathrm{H}}t - K_{\mathrm{H}}z\right)\cos\left(\Omega_{\mathrm{L}}t - K_{\mathrm{L}}z - \theta\right) \tag{10}$$

with the values listed in the table in the Supplementary Materials section III. The final signal is then renormalized by renorm. = max(signal) to be bounded between -1 and 1, so that the final radius is not exceeding $a_0 \pm \phi$:

$$a_{\mathrm{segmentation}}(t,z) = a_0\left(1 + \phi\,\mathrm{signal}(t,z)/\mathrm{renorm.}\right). \tag{11}$$

Changing the $\Gamma$ coefficients, we can obtain a straight tube (all coefficients are 0) or peristalsis (only the high-frequency wave coefficient is non-zero, see Supplementary Materials section III). Therefore, this is a general pattern of motility for the gut.

This pattern is still periodical in time, with periodicity given by the reciprocal of the greatest common denominator ($g.d.c$) of the frequencies $\Omega_{\mathrm{H}}, \Omega_{\mathrm{L}}, \Omega_{\mathrm{L}} + \Omega_{\mathrm{H}}$, and $\Omega_{\mathrm{L}} - \Omega_{\mathrm{H}}$. In fact, the signal can be rewritten as

$$\begin{aligned}
\mathrm{signal}(t,z) &= \Gamma_{\mathrm{H}}\sin\left(\Omega_{\mathrm{H}}t - K_{\mathrm{H}}z\right) + \Gamma_{\mathrm{L}}\sin\left(\Omega_{\mathrm{L}}t - K_{\mathrm{L}}z\right) + \Gamma_{\mathrm{P}}\sin\left(\Omega_{\mathrm{H}}t - K_{\mathrm{H}}z\right)\cos\left(\Omega_{\mathrm{L}}t - K_{\mathrm{L}}z - \theta\right) & (12)\\
&= \Gamma_{\mathrm{H}}\sin\left(\Omega_{\mathrm{H}}t - K_{\mathrm{H}}z\right) + \Gamma_{\mathrm{L}}\sin\left(\Omega_{\mathrm{L}}t - K_{\mathrm{L}}z\right) \\
&\quad + \frac{\Gamma_{\mathrm{P}}}{2}\left[\sin\left(\left(\Omega_{\mathrm{H}} - \Omega_{\mathrm{L}}\right)t - \left(K_{\mathrm{H}} - K_{\mathrm{L}}\right)z + \theta\right) + \sin\left(\left(\Omega_{\mathrm{H}} + \Omega_{\mathrm{L}}\right)t - \left(K_{\mathrm{H}} + K_{\mathrm{L}}\right)z - \theta\right)\right]. & (13)
\end{aligned}$$

#### II. FLOW CALCULATIONS

Given the motility pattern, which is represented by the radius $a(t,z)$ dynamics in time and space, the cross-sectional averaged flow velocity $U(t,z)$ can be calculated from conservation of volume [2]:

$$\frac{\partial\left(a^2 U\right)}{\partial z} = -\frac{\partial a^2}{\partial t} \tag{14}$$

We impose pressure boundaries conditions; practically, the tube is open for the fluid to go out and in. Then the condition for conservation of volume reads [3]:

$$U(t,z=0) = -\frac{1}{8}\frac{1}{a(t,z=0)^2}\left(\frac{\frac{\Delta p}{\mu} - 8\int_0^L a^{-4}(t,z_2)\int_0^{z_2}\frac{\partial a^2(z1,t)}{\partial t}\mathrm{d}z_1\mathrm{d}z_2}{\int_0^L a^{-4}(t,z_1)\mathrm{d}z_1}\right) \tag{15}$$

where the first part is the flow due to an external pressure drop $\Delta p$ (with $\mu$ being the fluid viscosity), and the second part is the pressure drop generated by the contracting tube walls. When considering peristalsis and segmentation, we impose this fixed external pressure drop to zero, so that the flow generated is only due to the motility pattern. For a straight tube instead, we have no contribution from the moving walls since $a(t,z) = a_0$ and we impose a fixed negative external pressure drop.

We define the equivalent average flow velocity over a period of contraction as $\langle U\rangle$. This is the flow velocity that a straight tube would have if we impose that the straight tube has the same period-average volumetric flow of the motility case $U(t,z)a(t,z)^2$:

$$U_{\text{straight}} := \frac{\frac{1}{T}\int_0^{\mathrm{T}} a(t,z)^2 U(t,z)\mathrm{d}t}{a_0^2}, \tag{16}$$

with $T$ the contraction time period. The period-averaged volumetric flow $\frac{1}{T}\int_0^{\mathrm{T}} a(t,z)^2 U(t,z)\mathrm{d}t$ oscillates in space around a well defined value due to the motility patterns space-dependence; therefore, we take also the spatial mean:

$$\langle U\rangle = \frac{1}{L}\int_0^L U_{\text{straight}}\mathrm{d}z \tag{17}$$

where $L$ is the tube length.

Note that our definition of equivalent average flow velocity is different from the period-averaged velocity:

$$\langle U\rangle \neq \frac{1}{T}\int_0^{\mathrm{T}} U(t,z)\mathrm{d}t \tag{18}$$

and therefore the equivalent average flow velocity can be higher than the motility wave velocity $\Omega/K$. Indeed, the important quantity for transport is the volumetric flow and not the instantaneous flow; therefore we use for comparison between patterns the equivalent velocity and not just the period-averaged velocity.

#### III. PARAMETERS

| Variable | Symbol | Value | Comments |
|---|---|---|---|
| Length of tube | $L$ | 0.33 m | murine S.I. length [4] |
| Radius | $a$ | 0.5 mm | Taylor limit, murine S.I. lumen is 0.77 mm [5] |
| occlusion | $\phi$ | 10 % | Taylor limit, human occlusion is 73% [6] |
| High-frequency wave frequency | $\Omega_{\mathrm{H}}$ | $2\pi \cdot 0.83$ Hz | high-frequency/peristalsis see Supp. section I [1] |
| High-frequency wave wavelength | $K_{\mathrm{H}}$ | $2\pi \cdot 98.0\ \mathrm{m}^{-1}$ | high-frequency/peristalsis see Supp. section I [1] |
| Low-frequency wave frequency | $\Omega_{\mathrm{L}}$ | $2\pi \cdot 0.13$ Hz | Modulating low-freq./segmentation see Supp. section I [1] |
| Low-frequency wave wavelength | $K_{\mathrm{L}}$ | $2\pi \cdot 190.5\ \mathrm{m}^{-1}$ | Modulating low-freq./segmentation see Supp. section I [1] |
| High-frequency wave amplitude | $\Gamma_{\mathrm{H}}$ | 0.78 | Amplitude of the high-frequency wave see Supp. sectionI [1] |
| Low-frequency wave amplitude | $\Gamma_{\mathrm{L}}$ | 1 | Amplitude of the low-frequency wave see Supp. section I [1] |
| Modulating wave amplitude | $\Gamma_{\mathrm{P}}$ | 0.48 | Amplitude of the modulating wave see Supp. section I [1] |
| Phase shift | $\theta$ | $2\pi \cdot 0.3$ | Checkerboard pattern requirement, see Supp. section I [1] |
| Viscosity | $\mu$ | 0.001 Pa s | Water viscosity [7] |
| Fixed pressure drop ($a = a(z,t)$) | $\delta P$ | 0 Pa | Pure pumping due to motility see Supp. section II |
| Fixed pressure drop ($a = a_0$) | $\delta P$ | Variable | Fixed pressure drop for straight tube see Supp. section II |
| Diffusion coefficient | $k$ | $2 \cdot 10^{-9}\ \mathrm{m}^2\mathrm{s}^{-1}$ | Molecule of 1 Å, nutrient and motile bacteria |
| absorption rate | $\tilde{\gamma}$ | $1 \cdot 10^{-6}\ \mathrm{m\ s}^{-1}$ | [7, 8] |
| Nutrients at inlet | $N_0$ | $100\ \mathrm{mol\ m}^{-3}$ | [7] |
| Bacteria at inlet | $B_0$ | $1.7 \cdot 10^{-14}\ \mathrm{mol\ m}^{-3}$ | ($1 \cdot 10^7$ cells $\mathrm{L}^{-1}$ ) [7] |
| Nutrients cutoff | $\tilde{N}$ | $20\ \mathrm{mol\ m}^{-3}$ | (0.02 mol $\mathrm{L}^{-1}$ ) [7] |
| Nutrient consumption rate | $\alpha_{BN}$ | $2.8 \cdot 10^7\ \mathrm{s}^{-1}$ | ($4.6 \cdot 10^{-17}$ mol cells$^{-1}$ s$^{-1}$ ) [7] |
| Bacteria growth rate | $\alpha_{\text{growth}}$ | $2 \cdot 10^{-3}\ \mathrm{s}^{-1}$ | [7] |
| Timestep of integration | $\Delta t$ | $7 \cdot 10^{-4}$ s | Numerical stability, see Supp. section V |
| Spacestep of integration | $\Delta z$ | $4 \cdot 10^{-4}$ m | Numerical stability, see Supp. section V |
| Gaussian width | | 30 $\Delta z$ | Width of the initial nutrient Gaussian for Fig. 2 of main |

#### IV. TAYLOR EXPANSION CALCULATION

##### A. Premises

In this section, we want to obtain the Taylor expansion equation for the nutrient cross-sectional averaged concentration $N$ with moving absorbing walls and bacterial consumption. First, we derive it without the term of bacterial consumption; then, this term is added in Section IV G. The bacterial equation is similar to the nutrients one, as we further discuss in the different sections. We base our derivation on Marbach's work using the Manifold method [2], but we extend the equations to include moving boundaries, nutrient absorption at the moving walls, and a Monod term.

In synthesis, we want to obtain an equation for the cross-sectional averaged concentration, which can be obtained supposing that the dynamics is dominated by the longitudinal dimension and that everything is equilibrated and well mixed radially at faster time scales. Mathematically speaking, we need to impose that diffusion time scales are faster than the advective one $\epsilon := \frac{Ua^2}{kL} < 1$, the so-called Taylor limit [2]. We also assume to have a small dimensionless absorption constant $\gamma_{\text{adim}} := \gamma a(t,z)/k$ of the same order as $\epsilon$, similarly to Marbach's work [2]. We do not necessarily have to choose $\gamma_{\text{adim}}$ and $\epsilon$ of the same order, and in principle, the two limits should be considered separately. However, since both terms are small, this assumption is safe [2].

##### B. The 2D equation

First, we start from the two dimensional advection-diffusion equation for the concentration $n = n(t,z,r)$, which depends on the longitudinal space coordinate $z$, the radial coordinate $r$, and time $t$. We consider moving boundaries with radius depending on space and time $a = a(t,z)$. The equation becomes

$$\frac{\partial n}{\partial t} = -u\frac{\partial n}{\partial z} - v\frac{\partial n}{\partial r} + k\frac{\partial^2 n}{\partial z^2} + k\frac{1}{r}\frac{\partial}{\partial r}\left(r\frac{\partial n}{\partial r}\right). \tag{19}$$

Where $u = u(t,z,r)$ and $v = v(t,z,r)$ are the longitudinal and radial velocity, respectively.

##### C. Boundary conditions

The nutrient absorption at the boundaries is written as the following condition:

$$\left(\frac{k}{a}\frac{\partial n}{\partial \eta} - k\frac{\partial a}{\partial z}\frac{\partial n}{\partial z}\right)|_{\eta=1} = -\gamma n|_{\eta=1} \tag{20}$$

where $\eta := r/a$ is the dimensionless radial variable. This boundary condition is derived as follows.

Equating the nutrient flux at the wall, we obtain [2]:

$$(k\nabla n)_{\perp,\,\eta=1} = -\gamma n|_{\eta=1} \tag{21}$$

where $(k\nabla n)_{\perp,\,\eta=1}$ is the perpendicular diffusive flux, and $-\gamma n|_{\eta=1}$ are the absorbed molecules at the wall. The perpendicular unit vector at the wall is $\hat{n}_\perp \simeq \hat{r} - \frac{\mathrm{d}a}{\mathrm{d}z}\hat{z}$, thanks to the lubrication assumption $\mathrm{d}a/\mathrm{d}z << 1$ [2], with $\hat{r}$ the unit radial vector; it also holds, for the same approximation, that $|\hat{n}_\perp| \simeq 1$. Using this vector we obtain:

$$(k\nabla n)_{\perp,\,\eta=1} = (k\nabla n\cdot\hat{n}_\perp)|_{\eta=1} = \left[k\left(\frac{\partial n}{\partial r}\hat{r} + \frac{\partial n}{\partial z}\hat{z}\right)\cdot\left(\hat{r} - \frac{\mathrm{d}a}{\mathrm{d}z}\hat{z}\right)\right]_{\eta=1} = k\left[\frac{\partial n}{\partial r} - \frac{\partial n}{\partial z}\frac{\mathrm{d}a}{\mathrm{d}z}\right]_{\eta=1} \equiv -\gamma n|_{\eta=1}, \tag{22}$$

and finally, using $\eta = r/a$ at first order we obtain Supplementary Eq. 20.

##### D. The invariant manifold method

We aim to write the equation for the cross-section averaged concentration defined as:

$$N(t,z) = \frac{1}{\text{area}}\int_{\text{area}} n\,\mathrm{d}S = \frac{1}{\pi a^2}\int_0^{2\pi}\mathrm{d}\phi\int_0^a n\,r\,\mathrm{d}r = 2\int_0^1 n\,\eta\,\mathrm{d}\eta. \tag{23}$$

To obtain the Taylor expansion equation for $N$, we will use the invariant manyfold method [9–13], which is reviewed and extended by Marbach [2].

We first rewrite the Supplementary Eq. 19 using the adimensional variables $\epsilon$, $\beta := k^2/(U^2a^2)$, with $U(t,z) = 2\int_0^1 u\,\eta$ , $r' := r/a$, $z' := z/L$, $t' := t/(L/U)$, $u' := u/U$ and $v' := v/(Ua/L)$. The equation becomes [2]:

$$\frac{1}{\epsilon}\mathcal{L}n = \frac{\partial n}{\partial t'} + u'\frac{\partial n}{\partial z'} + v'\frac{\partial n}{\partial r'} - \epsilon\beta\frac{\partial^2 n}{\partial z'^2} \tag{24}$$

where we also defined the operator:

$$\mathcal{L}n := \frac{1}{r'}\frac{\partial}{\partial r'}\left(r'\frac{\partial n}{\partial r'}\right) \tag{25}$$

According to the manifold expansion method, we write the derivative of the cross-section averaged concentration $\mathrm{d}N/\mathrm{d}t'$ and of the concentration $n$ as an expansion in $\epsilon$. With $\epsilon << 1$ (Taylor limit), we obtain an accurate description of the concentration with few orders of the expansion [2]. The expansions are

$$\frac{\partial N}{\partial t'} = G'^{(1)}\left[N, \frac{\partial N}{\partial z}\right] + \epsilon\, G'^{(2)}\left[N, \frac{\partial N}{\partial z'}, \frac{\partial^2 N}{\partial z'^2}\right] + ... \tag{26}$$

and

$$n = V'^{(0)}\left[r', t'; N\right] + \epsilon\, V'^{(1)}\left[r', t'; N, \frac{\partial N}{\partial z'}\right] + \epsilon^2\, V'^{(2)}\left[r', t'; N, \frac{\partial N}{\partial z'}, \frac{\partial^2 N}{\partial z'^2}\right] + ... \tag{27}$$

Now we can substitute these expansions back in the Supplementary Eq. 24, keeping in mind that when taking the derivative with respect to time, all the $V$ terms depend directly on $t'$ but also depend on the time through $N$, $\partial N/\partial z'$,... and similarly with the derivative with respect to space [2]. After substituting, we can compare the left side and right side with the same order of $\epsilon$. We thus get:

$$\mathcal{L}V'^{(0)} = 0 \tag{28}$$

$$\mathcal{L}V'^{(1)} = \frac{\partial V'^{(0)}}{\partial t'} + \frac{\partial V'^{(0)}}{\partial C}G'^{(1)} + u'\frac{\partial V'^{(0)}}{\partial z'} + v'\frac{\partial V'^{(0)}}{\partial r'} \tag{29}$$

$$\mathcal{L}V'^{(2)} = \frac{\partial V'^{(1)}}{\partial t'} + \frac{\partial V'^{(1)}}{\partial C}G'^{(1)} + \frac{\partial V'^{(1)}}{\partial\left(\partial C/\partial z'\right)}\frac{\partial G'^{(1)}}{\partial z'} + \frac{\partial V'^{(0)}}{\partial C}G'^{(2)} + u'\frac{\partial V'^{(1)}}{\partial z'} + v'\frac{\partial V'^{(1)}}{\partial r'} - \beta\frac{\partial^2 V'^{(0)}}{\partial z'^2} \tag{30}$$

Higher orders are not considered in the present paper but can be found in Marbach [2].

Substituting the adimensional variables with the corresponding dimensional variable (apart from the radius which will be kept as $\eta := r/a$), and considering at first order $\frac{\partial}{\partial r} \simeq \frac{1}{a}\frac{\partial}{\partial \eta}$, we obtain the equations we need to solve to write the Taylor expansion for $N$:

$$\boxed{\begin{aligned}
&\frac{1}{\eta}\frac{\partial}{\partial\eta}\left(\eta\frac{\partial V^{(0)}}{\partial\eta}\right) = 0 &(31)\\
&\frac{k}{a^2}\frac{1}{\eta}\frac{\partial}{\partial\eta}\left(\eta\frac{\partial V^{(1)}}{\partial\eta}\right) = \frac{\partial V^{(0)}}{\partial t} + \frac{\partial V^{(0)}}{\partial N}G^{(1)} + u\frac{\partial V^{(0)}}{\partial z} + v\frac{1}{a}\frac{\partial V^{(0)}}{\partial\eta} &(32)\\
&\frac{k}{a^2}\frac{1}{\eta}\frac{\partial}{\partial\eta}\left(\eta\frac{\partial V^{(2)}}{\partial\eta}\right) = \frac{\partial V^{(1)}}{\partial t} + \frac{\partial V^{(1)}}{\partial N}G^{(1)} + \frac{\partial V^{(1)}}{\partial\left(\partial N/\partial z\right)}\frac{\partial G^{(1)}}{\partial z} + \frac{\partial V^{(0)}}{\partial N}G^{(2)} + u\frac{\partial V^{(1)}}{\partial z} + v\frac{1}{a}\frac{\partial V^{(1)}}{\partial\eta} +\\
&\qquad\qquad - k\frac{\partial^2 V^{(0)}}{\partial z^2} &(33)
\end{aligned}}$$

where the expansion in dimensional variables now is:

$$\boxed{\frac{\partial N}{\partial t} = G^{(1)} + G^{(2)} + ...} \tag{34}$$

and

$$\boxed{n = V^{(0)} + V^{(1)} + V^{(2)} + ...} \tag{35}$$

keeping in mind that $V^{(0)} = V'^{(0)}$, $V^{(1)} = \epsilon V'^{(1)}$, ..., and $G^{(1)} = G'^{(1)}$, $G^{(2)} = \epsilon G'^{(2)}$, ...

We now have to impose the boundary condition, which becomes, using the same expansions of Supplementary Eq. 26 and Supplementary Eq. 27 in Supplementary Eq. 20, and comparing the correct orders in $\epsilon$:

$$\frac{k}{a}\frac{\partial V^{(0)}}{\partial \eta}|_{\eta=1} = 0 \tag{36}$$

$$\frac{k}{a}\frac{\partial V^{(1)}}{\partial \eta}|_{\eta=1} + \gamma V^{(0)}|_{\eta=1} = 0 \tag{37}$$

$$\frac{k}{a}\frac{\partial V^{(2)}}{\partial \eta}|_{\eta=1} - k\frac{\partial a}{\partial z}\frac{\partial V^{(0)}}{\partial z}|_{\eta=1} + \gamma V^{(1)}|_{\eta=1} = 0 \tag{38}$$

##### E. Additional relations

To continue in our derivation, we assume we are in the lubrication limit and that there is a Poiseuille flow [2]:

$$u = 2U(1-\eta^2) \tag{39}$$

$$v = 2\frac{\partial a}{\partial z}U(\eta - \eta^3) + \frac{\partial a}{\partial t}\eta(2-\eta^2). \tag{40}$$

where $U$ is the cross-sectional averaged longitudinal flow.

Additional relations we used to solve the equations and find the expansion orders are the following [2]:

$$\int_0^1 2V^{(0)}\eta \mathrm{d}\eta = N, \; \int_0^1 2V^{(i>0)}\eta \mathrm{d}\eta = 0 \tag{41}$$

This derives from the fact that we can just assume that all the contributions come from the first term in the expansion:

$$N := \int_0^1 2\eta n \; \mathrm{d}\eta = \int_0^1 2\eta \left(V^{(0)} + V^{(1)} + V^{(2)} + ...\right) \mathrm{d}\eta. \tag{42}$$

We can also assume [2] that:

$$\frac{\partial V^{(i)}}{\partial \eta}|_{\eta=0} = finite \tag{43}$$

All these relations are used to find the expansion terms for $n$ and $\mathrm{d}N/\mathrm{d}t$, and thus to obtain a Taylor equation for $N$. We will solve one order at a time.

##### F. Derivation

###### 1. Zeroth order

We want to solve the zeroth order Supplementary Eq. 31 with boundary Supplementary Eq. 36:

$$\frac{1}{\eta}\frac{\partial}{\partial \eta}\left(\eta\frac{\partial V^{(0)}}{\partial \eta}\right) = 0 \tag{44}$$

$$\frac{\partial V^{(0)}}{\partial \eta}|_{\eta=1} = 0 \tag{45}$$

Integrating once in $\eta$, $\eta\frac{\partial V^{(0)}}{\partial\eta} + const.' = 0$. To calculate the constant, lets consider it in $\eta = 1$; using the boundary, $const.' = 0$. Thus we get that $V^{(0)} = const.$. Using the additional relations Supplementary Eq. 41, we have $N = \int_0^1 2V^{(0)}\eta d\eta = V^{(0)}\int_0^1 2\eta d\eta = V^{(0)}$ so we get

$$\boxed{V^{(0)} = N} \tag{46}$$

*2. First order*

We solve here the first order Supplementary Eq. 32 with boundary Supplementary Eq. 37:

$$\frac{k}{a^2}\frac{1}{\eta}\frac{\partial}{\partial\eta}\left(\eta\frac{\partial V^{(1)}}{\partial\eta}\right) = \frac{\partial V^{(0)}}{\partial t} + \frac{\partial V^{(0)}}{\partial N}G^{(1)} + u\frac{\partial V^{(0)}}{\partial z} + v\frac{1}{a}\frac{\partial V^{(0)}}{\partial\eta} \tag{47}$$

$$\frac{k}{a}\frac{\partial V^{(1)}}{\partial\eta}|_{\eta=1} + \gamma V^{(0)}|_{\eta=1} = 0. \tag{48}$$

using the expression for $u$ Supplementary Eq. 39 and $V_0$ Supplementary Eq. 46 (from which it follows that $\frac{1}{a}\frac{\partial V^{(0)}}{\partial\eta} = \frac{1}{a}\frac{\partial N}{\partial\eta} = 0$ so that the term in $v$ drops out from the equation), similarly from solution for the zeroth order, we integrate once for $\eta$. The constant of integration is 0 since we impose that the derivative in $\eta$ of $V^{(1)}$ is finite for $\eta = 0$ (Supplementary Eq. 43). Then, we use the boundary condition at $\eta = 1$, and we find that $G^{(1)}$ is the same as found by Marbach for a straight tube with absorbing walls [2]:

$$\boxed{G^{(1)} = -U\frac{\partial N}{\partial z} - 2N\frac{\gamma}{a}} \tag{49}$$

Using now $G^{(1)}$ in the integral form of $V^{(1)}$, and using the fact that the cross-sectional average of $V^{(1)}$ has to vanish (Supplementary Eq. 41), we obtain again as in a straight tube with absorbing walls [2]:

$$\boxed{V^{(1)} = \frac{1}{4}\left(1 - 2\eta^2\right)\gamma\frac{a}{k}N + \frac{\partial N}{\partial z}\frac{Ua^2}{24k}\left(-2 + 6\eta^2 - 3\eta^4\right).} \tag{50}$$

Until the first order in the expansion, there are no effects of the moving boundaries but only of the absorbing ones, consistently with Marbach [2].

*3. Second order*

We solve here the second order Supplementary Eq. 33 with boundary Supplementary Eq. 38:

$$\frac{k}{a^2}\frac{1}{\eta}\frac{\partial}{\partial\eta}\left(\eta\frac{\partial V^{(2)}}{\partial\eta}\right) = \frac{\partial V^{(1)}}{\partial t} + \frac{\partial V^{(1)}}{\partial N}G^{(1)} + \frac{\partial V^{(1)}}{\partial\left(\partial N/\partial z\right)}\frac{\partial G^{(1)}}{\partial z} + \frac{\partial V^{(0)}}{\partial N}G^{(2)} + u\frac{\partial V^{(1)}}{\partial z} + v\frac{\partial V^{(1)}}{\partial r} - k\frac{\partial^2 V^{(0)}}{\partial z^2} \tag{51}$$

$$\frac{k}{a}\frac{\partial V^{(2)}}{\partial\eta}|_{\eta=1} - k\frac{\partial a}{\partial z}\frac{\partial V^{(0)}}{\partial z}|_{\eta=1} + \gamma V^{(1)}|_{\eta=1} = 0. \tag{52}$$

To proceed, we calculate the derivative of $V^{(1)}$ in $t$, $z$ $\eta$ (keeping in mind that $\partial V^{(1)}/\partial z \neq 0$ and that it depends also on $a$ and $\eta$). We then substitute the derivatives and $G^{(1)}$ as well into the second order equation, obtaining:

$$\begin{aligned}\frac{\partial}{\partial\eta}\left(\eta\frac{\partial V^{(2)}}{\partial\eta}\right) = N\left[\frac{a^2}{k^2}\gamma^2\left(\eta^3 - \frac{\eta}{2}\right) + a^2\frac{\partial a}{\partial t}\frac{\gamma}{4k^2}\left(\eta - 6\eta^3 + 4\eta^5\right) + \frac{a^2}{2}\frac{\partial a}{\partial z}\frac{\gamma}{k^2}U\left(\eta - 3\eta^3 + 2\eta^5\right)\right] +\\ \frac{\partial N}{\partial z}\left[\gamma\frac{U}{4k^2}a^3\left(\eta - 4\eta^3 + 4\eta^5\right) + \frac{\partial a}{\partial t}\frac{a^3U}{12k^2}\left(2\eta - 4\eta^3 + 3\eta^5\right)\right]\\ + \frac{\partial^2 N}{\partial z^2}\left[-a^2\eta + \frac{a^4}{12k^2}U^2\left(-2\eta + 8\eta^3 - 9\eta^5 + 3\eta^7\right)\right] + G^{(2)}\left[\frac{a^2}{k}\eta\right].\end{aligned}$$

Our aim here is to determine $G^{(2)}$. So we integrate the last equation once in $\eta$, keeping in mind that the constant of this integration is 0 because we impose that $\partial V^{(2)}/\partial\eta$ is finite in $\eta = 0$ (Supplementary Eq. 43). Substituting $V^{(1)}$ in the boundary condition, and applying the boundary at $\eta = 1$ on the integrated equation, we find for $G^{(2)}$:

$$\boxed{G^{(2)} = N\left[\frac{\partial a}{\partial t}\frac{\gamma}{6k} - \frac{\partial a}{\partial z}U\frac{\gamma}{12k} + \frac{\gamma^2}{2k}\right] + \frac{\partial N}{\partial z}\left[-\gamma U\frac{a}{6k} + \frac{\partial a}{\partial z}\frac{2k}{a} - \frac{\partial a}{\partial t}U\frac{a}{12k}\right] + \frac{\partial^2 N}{\partial z}\left[k + U^2\frac{a^2}{48k}\right].} \tag{53}$$

Then, using the expression for $G^{(1)}$ (Supplementary Eq. 49) and $G^{(2)}$ (Supplementary Eq. 53), we find at second order using Supplementary Eq. 34:

$$\frac{\partial N}{\partial t} = G^{(1)}+G^{(2)} = -2\frac{\gamma}{a}\left(1 - \frac{\gamma a}{4k} - \frac{a}{12k}\frac{\partial a}{\partial t} + \frac{Ua}{24k}\frac{\partial a}{\partial z}\right)N - U\left(1 + \gamma\frac{a}{6k} - \frac{2k}{aU}\frac{\partial a}{\partial z} + \frac{a}{12k}\frac{\partial a}{\partial t}\right)\frac{\partial N}{\partial z} + k\left(1 + \frac{a^2U^2}{48k^2}\right)\frac{\partial^2 N}{\partial z^2}, \tag{54}$$

which using the relation $\frac{\partial a^2}{\partial t} = 2a\frac{\partial a}{\partial t}$ and similarly in $z$, becomes equation 2 of the main manuscript with equations from the main manuscript 3, 4, and 5 without the bacterial Monod term. Note that if $a = a_0$ we obtain the equation for a straight tube with absorbing walls [2]; if we set $\gamma = 0$ we obtain the equation for a tube with moving walls but no absorption, which is the basis of the equation we use for the bacteria [2]; and setting both conditions gives the basic Taylor dispersion equation for a straight tube without absorption [2].

##### G. Adding a Monod term

In our work, bacterial and nutrient equations are coupled through a Monod term [7, 14, 15]. This term is a bulk sink term for the nutrient equation since nutrients are consumed in the bulk and not at the boundary as the wall absorption term; instead, it is a source bulk term for bacteria, since bacteria growth in the bulk in our model and we do not consider adhesion to the walls.

Since $B$ and $N$ are radially well mixed under our hypothesis, we add to the Taylor expansions the Monod terms, giving an effective description of the bulk loss of nutrients due to bacteria and the bacterial growth due to nutrients. Simplifying the equations from Ishikawa [7] by neglecting oxygen and reducing our bacteria to one species only, we add to the nutrient equation the term $-\left(\alpha_{\mathrm{BN}}B\right)\frac{N}{N+\bar{N}}$ and to the bacterial equation the term $+\left(\alpha_{\mathrm{g}}\frac{N}{N+\bar{N}}\right)B$.

#### V. NUMERICAL DETAILS

We used an Euler forward explicit method, with central difference derivatives. The nutrient equation without the bacteria term, once discretized, becomes:

$$\begin{aligned} N_m^{j+1} = N_m^j &- 2\Delta t\frac{k}{\left(a^2\right)_m^j}\gamma_m^j\left(1 - \frac{\gamma_m^j}{4} - \frac{1}{24k}\left(\frac{\partial a^2}{\partial t}\right)_m^j + \frac{U_m^j}{48k}\frac{\partial\left(a^2\right)_m^j}{\partial z}\right)N_m^j + \\ &-\Delta t\left(U_m^j + U_m^j\frac{\gamma_m^j}{6} - \frac{k}{\left(a^2\right)_m^j}\left(\frac{\partial a^2}{\partial z}\right)_m^j + \frac{U_m^j}{24k}\left(\frac{\partial a^2}{\partial t}\right)_m^j\right)0.5\frac{N_{m+1}^j - N_{m-1}}{\Delta z} + \\ &+\Delta t\, k\left(1 + \frac{\left(a^2\right)_m^j\left(U^2\right)_m^j}{48k^2}\right)\frac{N_{m+1}^j + N_{m-1}^j - 2N_m^j}{\Delta z^2} \end{aligned} \tag{55}$$

Stability conditions for an advective-diffusive equation are the Von Neumman condition and the Courant–Friedrichs–Levy condition [16]:

$$k_{\mathrm{eff}}\frac{\Delta t}{\Delta z^2} < 0.5 \tag{56}$$

$$0.5U_{\mathrm{eff}}\frac{\Delta t}{\Delta z} < 1 \tag{57}$$

For a diffusion equation with absorption, applying the Von Neumann stability method, we obtain also the additional condition:

$$\gamma_{\text{eff}} \Delta t < 1 \tag{58}$$

The simulation is written in Fortran 90.

#### VI. FLUX AND INLET/OUTLET BOUNDARY CONDITIONS

##### A. Flux

The flux is calculated without the nutrient absorption term and sink/growth terms, for a tube with moving walls. We can write the Taylor expansion equation in the form [2]:

$$\frac{\partial \rho}{\partial t} + \frac{\partial J_{\text{N}}}{\partial z} = 0 \tag{59}$$

with $\rho = Na^2$. The current thorough a section of the tube (mol/s) is defined as [2]:

$$J_{\text{N}} = U\rho - k_{\text{eff}} \frac{\partial \rho}{\partial z} + \rho a^2 k_{\text{eff}} \frac{\partial a^2}{\partial z} = Na^2 U - k_{\text{eff}} a^2 \frac{\partial N}{\partial z} \tag{60}$$

which is valid in general for any pattern $a(z, t)$. The current at the boundary then becomes

$$J_{\text{N}}\big|_{\text{boundary}} = \left( a^2 NU - a^2 k_{\text{eff}} \frac{\partial N}{\partial z} \right) \Bigg|_{\text{boundary}} \tag{61}$$

and technically the flux is given by $J_{\text{N}}/area$, which for a straight tube is $\left(NU - 2k_{\text{eff}} \frac{\partial N}{\partial z}\right)\Big|_{\text{boundary}}$. In the text, we call flux the current $J_{\text{N}}$.

Considering that $influx = outflux + walluoptake$, we estimate the error when adding the absorption term in the nutrient equation to be around 2%.

##### B. Boundary conditions

Here we define the inlet/outlet boundary conditions used through the paper: no flux boundary, upheld boundary and free outflow boundary.

- no flux boundary: nothing leaves the tube at the boundary

$$NU^2 - k_{\text{eff}} \frac{\partial N}{\partial z}\Bigg|_{\text{boundary}} = 0 \tag{62}$$

- free outflow boundary: molecules leave the tube due to the flow, but not by diffusion

$$\frac{\partial N}{\partial z}\Bigg|_{\text{boundary}} = 0 \tag{63}$$

- upheld concentration: there is a reservoir at the boundary

$$N\Bigg|_{\text{boundary}} = N_0 = const. \tag{64}$$

#### VII. ABSORPTION RATE

We want to calculate the rate of nutrient absorption through the wall, thus the number of mols per second that pass through the wall boundary. We thus consider the flux perpendicular to the wall of the nutrient concentration $n = n(t, z, r)$

$$\Phi_{\mathrm{N}} := -\int_{\mathrm{Surf}} (k\nabla n)_{\perp} \mathrm{d}S \tag{65}$$

using the boundary condition $(k\nabla n)_{\perp} = -\gamma n|_{\eta=1}$, and knowing that the general integration on a cylindrical surface can be written as $\int_{\mathrm{Surf}} f \mathrm{d}S = 2\pi \int_0^L \mathrm{d}z a f|_{\eta=1}$, we obtain

$$\Phi_{\mathrm{N}} = 2\pi\gamma \int_0^L \mathrm{d}z a n|_{\eta=1} \simeq 2\pi\gamma \int_0^L \mathrm{d}z a(V_0 + V_1 + ...)|_{\eta=1} \tag{66}$$

where we used the Taylor expansion of $n$ till the first order. Substituting $V_0$ and $V_1$ at $\eta = 1$ we obtain finally

$$\Phi_{\mathrm{N}} = 2\pi k \int_0^L \mathrm{d}z \frac{\gamma a}{k} \left[ N\left(1 - \frac{\gamma a}{4k}\right) + \frac{\partial N}{\partial z}\frac{Ua^2}{24k} \right] := \int_0^L \mathrm{d}z \left[ AN + B\frac{\partial N}{\partial z} \right] \tag{67}$$

#### VIII. NUTRIENT GAUSSIAN EXPERIMENT - THEORETICAL SOLUTION

In a dye injection experiment, a finite amount of dye is locally released in the gut [17]. The number of molecules crossing the walls and thus being absorbed, as well as the out-flux, can be measured. Similarly, in our simulation, we study the advection-dispersion and absorption of an initial Gaussian nutrient distribution, which can freely exit the outlet. No more solute enters the tube during the experiment. The initial concentration is a Gaussian centered in $z_0 = L/2$ given by

$$N = N_0 \mathrm{e}^{-\frac{(z-z_0)^2}{2\sigma^2}}. \tag{68}$$

The total initial concentration in the tube is

$$N_{\mathrm{initial}} = \int_0^L N \mathrm{d}z = N_0 \sigma \sqrt{2\pi} \mathrm{erf}\left(\frac{L}{2\sqrt{2}\sigma}\right), \tag{69}$$

which for a straight tube is proportional to the initial amount of molecules in the tube $\pi a_0^2 N_{\mathrm{initial}}$, which is also the maximum number of molecules which can be absorbed. Therefore, $N_{\mathrm{initial}}$ it is used as normalization.

Integrating the Taylor equation for a straight infinite tube, we obtain the concentration as a function of time and space:

$$N = N_0 \frac{\sigma}{\sqrt{\sigma^2 + 2k_{\mathrm{eff}} t}} \mathrm{e}^{-\frac{(z - z_0 - U_{\mathrm{eff}} t)^2}{2(\sigma^2 + 2k_{\mathrm{eff}} t)}} \mathrm{e}^{-\gamma_{\mathrm{eff}} t} \tag{70}$$

The solution is a Gaussian, with the center moving along the tube with velocity $U$, with the width increased by diffusivity, and with a decay due to the absorption. Using the definition of absorption rate (Supplementary Materials section VII), we can calculate the theoretical absorption rate

$$\begin{aligned} \Phi_{\mathrm{dye}} = N_0 \sigma \frac{\exp(-\gamma_{\mathrm{eff}} t)}{2\sqrt{\sigma^2 + 2k_{\mathrm{eff}} t}} \cdot & \\ \left[ -\frac{B}{\sqrt{\sigma^2 + 2k_{\mathrm{eff}} t}} \left( \mathrm{e}^{-\frac{(tU_{\mathrm{eff}} + z_0)^2}{\sigma^2 + 2k_{\mathrm{eff}} t}} - \mathrm{e}^{-\frac{(-L + tU_{\mathrm{eff}} + z_0)^2}{\sigma^2 + 2k_{\mathrm{eff}} t}} \right) \right. & \\ \left. + A\sqrt{2\pi} \left( \mathrm{erf}\left[ \frac{tU_{\mathrm{eff}} + z_0}{\sqrt{2(\sigma^2 + 2k_{\mathrm{eff}} t)}} \right] - \mathrm{erf}\left[ \frac{tU_{\mathrm{eff}} + z_0 - L}{\sqrt{2(\sigma^2 + 2k_{\mathrm{eff}} t)}} \right] \right) \right] & \end{aligned} \tag{71}$$

with all the effective coefficients with suffix "eff" and the coefficient $A$ and $B$ from Supplementary Eq. 67 calculated for a straight tube with condition $a = a_0$, and $U = U(t, 0)_{a=a_0}$. To get the total number of molecules in the tube, we need to integrate this till infinite times, which is not analytically solvable. Therefore, we numerically integrate the theoretical absorption rate of Supplementary Eq. 71 as we do with the simulated absorption rate, and we normalize it by the theoretical $N_{\mathrm{initial}}$. The results are plotted in Fig. 2C of the main manuscript.

#### IX. THEORETICAL ABSORPTION TIME

To calculate the decay time of the absorption rate $\Phi_{\rm N}$, we do a crude approximation, which proves to be valid since the fit is in agreement with simulations. We assume a straight tube, filled uniformly with concentration $N_0$. We assume no flow, and we neglect the diffusive term as well. Thus we can consider $N$ to be a constant in $z$. The equation to be solved thus becomes

$$\frac{\partial N}{\partial t} = -\gamma_{\rm eff}\Big|_{\rm a=a_0} N =: -\frac{1}{\tau_{\rm abs}} N \tag{72}$$

which has the straightforward solution $N = N_0 \exp(-t/\tau_{\rm abs})$. Using the definition of the absorption rate Supplementary Eq. 67, and using $N$ constant in $z$, we obtain

$$\Phi_{\text{no U, no diff.}} = 2\pi a L \frac{N_0}{\tau_{\rm abs}} \exp(-t/\tau_{\rm abs}); \qquad \tau_{\rm abs} = \gamma_{\rm eff}^{-1}\Big|_{a=a_0} = \left(\frac{2\gamma}{a} - \frac{\gamma^2}{2k}\right)^{-1} \tag{73}$$

This value is independent in good approximation from the motility pattern. To confirm this, for the nutrient Gaussian experiment, we show the absorption rate curve as a function of time, for peristalsis 10% occlusion, segmentation 10% occlusion, a straight tube with the same mean fluid velocity as peristalsis, and a straight tube with no flow (see Fig. 1). All curves follow the same decay process $\tau_{\rm abs}$ before the solute starts leaving the tube. Differences in the absorption rate curve after the solute leaves the tube can be explained by Taylor-dispersion-induced differences in the out-flux. However, the total absorbed molecules do not depend on motility, but only on the mean velocities and thus on residence times.

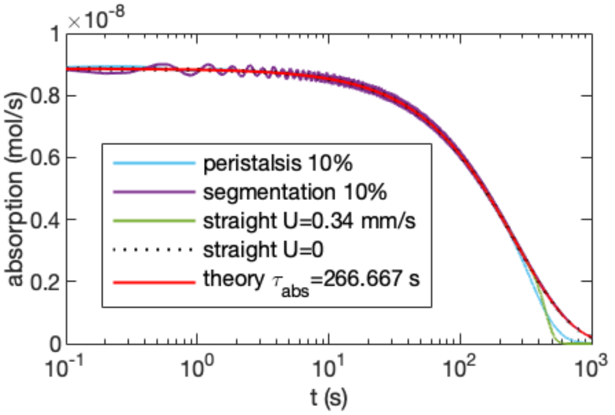


FIG. 1. **Absorption rate curve $\phi_{\rm N}$ as function of time for a nutrient Gaussian experiment**. Light blue 10% peristalsis, purple 10% segmentation, green straight tube with same velocity as peristalsis 10%, dotted line straight tube with no flow, red line exponential with decay time $\tau_{\rm abs} = \left(\frac{2\gamma}{a} - \frac{\gamma^2}{2k}\right)^{-1} = 266.667$ for $\gamma = 10^{-6}$ ms$^{-1}$.

#### X. STEADY-STATE FOR A STRAIGHT TUBE WITH AN UPHELD CONCENTRATION - THEORETICAL SOLUTION

##### A. Steady state equations

To calculate the steady-state for the nutrients, we impose the steady-state definition $\frac{\partial N}{\partial t} = 0$; we assume that bacteria do not influence the absorption of nutrients; we assume that diffusion can be neglected; we assume a straight tube; and an upheld concentration at the inlet $N(z = 0) = N_0$. Imposing all these conditions on Eq. 2 of the main manuscript, we solve and obtain:

$$N|_{\rm st.st.} = N_0 \exp\left[-\frac{\gamma_{\rm eff}}{U_{\rm eff}}|_{\rm a=a_0} z\right]. \tag{74}$$

Given that $\gamma_{\rm eff}|_{a=a_0} = 2\frac{\gamma}{a_0}\left(1-\frac{\gamma a_0}{4k}\right) = \tau_{\rm abs}^{-1}$, and that $U_{\rm eff}|_{a=a_0} = U\left(1+\frac{\gamma a_0}{6k}\right) \simeq U = L/\tau_{res}$, we can safely approximate the steady state solution with Eq. 7 from the main manuscript.

For the bacteria, we similarly assume a straight tube and we drop the diffusive term, while we retain the growth term depending on the nutrients. Imposing these conditions on Eq. 6 of the main manuscript and using Supplementary Eq. 74, we obtain:

$$B|_{\rm st.st.} = B_0\left(1+\frac{\bar{N}}{N_0}\right)^{\frac{\alpha_{\rm g}}{\gamma_{\rm eff}}\frac{U_{\rm eff}}{U}|_{a=a_0}}\left(\frac{\bar{N}}{N_0}+\exp\left(-\frac{\gamma_{\rm eff}}{U_{\rm eff}}|_{\rm a=a_0}z\right)\right)^{-\frac{\alpha_{\rm g}}{\gamma_{\rm eff}}\frac{U_{\rm eff}}{U}|_{a=a_0}} \tag{75}$$

which, with the same approximations made for the nutrients equations, we can safely approximate with Eq. 8 from the main manuscript.

##### B. Steady state absorption rate

Using Supplementary Eq. 67 and Supplementary Eq. 74, considering that $\frac{\partial N}{\partial z}|_{\rm st.st.} = -\frac{\gamma_{\rm eff}}{U_{\rm eff}}|_{1=1_0}N|_{\rm st.st.}$ the absorption rate becomes

$$\Phi_{\rm N}|_{\rm st.st.} = 2\pi k\int_0^L {\rm d}z\left[\frac{\gamma a_0}{k}\left(1-\frac{\gamma a_0}{4k}\right)-\frac{\gamma_{\rm eff}}{U_{\rm eff}}|_{a=a_0}\frac{Ua_0^2}{24k}\frac{\gamma a_0}{k}\right]N|_{\rm st.st.} \tag{76}$$

which, using Eq. 7 from the main manuscript and $\frac{\gamma_{\rm eff}}{U_{\rm eff}}|_{a=a_0} \simeq \frac{\tau_{\rm res}}{\tau_{\rm abs}L}$, and integrating along $z$ considering that for a straight tube the only dependence on the spatial variable is in the exponent in $N|_{\rm st.st.}$, we obtain:

$$\Phi_{\rm N}|_{\rm st.st.} = \frac{2\pi k\tau_{\rm abs}L}{\tau_{\rm res}}N_0\left[\frac{\gamma a_0}{k}\left(1-\frac{\gamma a_0}{4k}\right)-\frac{Ua_0^2}{24k}\frac{\gamma a_0}{k}\frac{\tau_{\rm res}}{L\tau_{\rm abs}}\right]\left(1-{\rm e}^{-\frac{\tau_{\rm res}}{\tau_{\rm abs}}}\right). \tag{77}$$

For visualization purposes in Fig. 3 of the main manuscript, we chose to normalize the absorption rate curves by the straight-tube infinite-velocity limit

$$\Phi_{\rm Uinf} = 2\pi kLN_0\frac{\gamma a_0}{k}(1-\frac{\gamma a_0}{4k})(1+\frac{\gamma a_0}{6k})(1+\frac{\gamma a_0}{12k})^{-1}. \tag{78}$$

This is calculated applying the limit of $U\to\infty$ to Supplementary Eq. 76, considering that the limit $\lim\limits_{U\to\infty} U\frac{\gamma_{\rm eff}}{U_{\rm eff}}|_{a=a_0}N_{\rm st.st.}$ is finite.

##### C. Efficiency definition for steady state

Since higher velocities absorb more molecules in absolute terms due to the higher influx, to quantify how good a tube is at absorbing we normalize the absorption rate $\Phi_{\rm N}$ by the influx, so that we measure the number of molecules which are not wasted by the flow compared to the incoming molecules. We define the steady-state efficiency as:

$$eff := 100\cdot\left.\frac{\Phi_{\rm N}}{J_{\rm N}|_0}\right|_{\rm st.st.}. \tag{79}$$

Efficiency can be higher than 100% because we used as normalization the influx of molecules calculated without nutrient absorption. The absorption terms in the equation would add corrections to the influx, providing the correct normalization. Thus, using the steady state solution Supplementary Eq. 74 we can calculate the theoretical efficiency which becomes

$$eff = 100\cdot\left(1-{\rm e}^{-\frac{\gamma_{\rm eff}}{U_{\rm eff}}|_{\rm a=a_0}L}\right)\left(1+\frac{\gamma a_0}{12k}\right)\left(1+\frac{k_{\rm eff}}{U}\frac{\gamma_{\rm eff}}{U_{\rm eff}}|_{\rm a=a_0}\right)^{-1} \tag{80}$$

and using again that $\gamma_{\rm eff}/U_{\rm eff}|_{a=a_0} \simeq \tau_{\rm res}/(\tau_{\rm abs}L)$ , we obtain

$$eff = 100\cdot\left(1-{\rm e}^{-\frac{\tau_{\rm res}}{\tau_{\rm abs}}}\right)\left(1+\frac{\gamma a_0}{12k}\right)\left(1+\frac{k_{\rm eff}\tau_{\rm res}}{LU\tau_{\rm abs}}\right)^{-1} \tag{81}$$

This solution is only valid for regimes of velocities within the Taylor limit but where the diffusive term can be ignored.

##### D. Proof of hypothesis

We used as a hypothesis to calculate the steady-state that bacteria consumption in the nutrient equation and diffusivity can be neglected. We showed good agreement in the main text in Fig. 3 of the main manuscript between straight-tube simulated data with bacterial terms and diffusivity, and our theory without these terms. Here, we show that even a simulation without a bacterial term is consistent with the simulation with the bacterial term, for what concerns efficiency. For a straight tube with a pressure drop of $\frac{\Delta P}{L} = -32$ Pa m$^{-1}$ and a corresponding velocity of 1 mm s$^{-1}$, we expect a theoretical efficiency value 70.3056% (calculated without bacteria and diffusive terms), we get from the simulations with bacteria consumption 70.7265%, and 70.7266% without bacteria consumption, confirming our hypothesis.

##### E. Bacterial growth definition and limit of validity

A good quantifier of bacterial dynamics is the number of bacteria in the tube at steady-state, normalized by the number of bacteria in a tube if the tube will be completely uniformly filled by the upheld concentration $B_0$:

$$\frac{b}{b_0} := \frac{\pi a^2 \int_0^L B \mathrm{d}z}{\pi a^2 L B_0} \tag{82}$$

This value shows growth if higher than 1. This value can not be written analytically, thus we numerically integrate this value once the theoretical $B$ is known. This value has a limit of 1 for high velocities, thus for high velocities, there is no growth.

This steady-state solution is valid only if bacterial consumption can be neglected compared to the absorption by the gut, so only if $\gamma_{\text{eff}} N >> \alpha_{\text{BN}} B \frac{N}{N+\bar{N}}$. Using the steady-state solution as an approaching limit to this validity point, and introducing the typical timescales of the system, we get that the equation is valid if:

$$\frac{\tau_{\text{abs}}}{\tau_{\text{cons}}} \frac{B_0}{N_0} \frac{(1+\frac{\bar{N}}{N_0})^{\frac{\tau_{\text{abs}}}{\tau_{\text{growth}}}}}{(\frac{\bar{N}}{N_0} + \exp(-\frac{\tau_{\text{res}}}{\tau_{\text{abs}}}))^{\frac{\tau_{\text{abs}}}{\tau_{\text{growth}}}+1}} << 1 \tag{83}$$

with $\tau_{\text{cons}} := 1/\alpha_{\text{B,N}}$ the consumption rate of the bacteria over the nutrients. Note that the important value that should be small is not $\frac{\tau_{\text{abs}}}{\tau_{\text{cons}}}$ but $\frac{\tau_{\text{abs}}}{\tau_{\text{cons}}} B_0/N_0$, since $\alpha_{\text{BN}}$ is weighting how many nutrients a certain number of bacteria are consuming.

#### XI. ABSORPTION AND FLUSHING PHASE AND THEIR FEEDBACK MECHANISMS

##### A. The phases and the feedback mechanisms

In the main Fig. 4, we provide an example of dynamics of the absorption phase, flushing phase, and of the feedback mechanisms involved in the switching between phases.

In the absorption (postprandial) phase, the meal is moved from the stomach to the small intestine. Then, the digestion of the meal starts, without a further income of nutrients or bacteria from the stomach. Therefore, we simulate the absorption phase as a tube with no nutrient influx or bacterial influx, with segmentation at 10% occlusion, with free bacterial and nutrients outflow. Initially, the tube is uniformly filled with bacterial concentration $B_0$ and with nutrient concentration $N_{\text{initial}}$. There are hints that nutrient presence can start this absorption phase thanks to nutrient-based feedback mechanisms [1, 18, 19], which would determine then the initial quantity of nutrients in the tube, $N_{\text{initial}}$.

The flushing phase happening during starvation is here represented with peristalsis at 10% occlusion, no influx of nutrients nor bacteria, and free outflow of nutrients and bacteria. The initial conditions are given by the final conditions of the absorption phase. To start the flushing phase, there are two possible mechanisms. If the tube is empty of nutrients, fast modes are observed to start [20]; we refer to this as a lack-of-nutrient feedback mechanism. Alternatively, the bacterial metabolites can start a fast flow phase as peristalsis [21]. We refer to this as bacterial threshold feedback, and for sake of simplicity in our simulation, peristalsis is started when bacteria reach a bacterial threshold $B_{\text{high}}$. Varying this threshold makes the absorption phase change in duration.

##### B. Timescales of the system

Here we summarize the timescales of the system and their meaning.

- $\tau_{\mathrm{abs}}$ absorption timescale. As shown in section IX, if nutrients are not leaving the tube due to advection, the nutrients in the tube can be described by $N = N_0 \exp(-t/\tau_{\mathrm{abs}})$.

- $\tau_{\mathrm{g}}$ bacterial growth rate.

- $\tau_{\mathrm{res}}^{\mathrm{per}}$, $\tau_{\mathrm{res}}^{\mathrm{seg}}$ respectively, peristalsis and segmentation residence times. These correspond to the time the solute spend in the tube, thus they give an idea of the timescale needed to empty the tube. They depend strongly on the flow velocity of the system.

- $T_{\mathrm{abs.\ phase}}$ the duration of the absorption phase, which is determined by the feedback mechanism type. If the lack-of-nutrient feedback is used, the phase is ended when the tube is emptied from the nutrients. The emptying timescale is represented by the residence time $\tau_{\mathrm{res}}^{\mathrm{seg}}$, therefore we obtain $T_{\mathrm{abs.\ phase}} \simeq \tau_{\mathrm{res}}^{\mathrm{seg}}$. To maximize absorption, slow flows need to be chosen so that $T_{\mathrm{abs.\ phase}} \simeq \tau_{\mathrm{res}}^{\mathrm{seg}} \gg \tau_{\mathrm{abs}}$. Instead, if the bacterial feedback mechanism is used, then the phase is stopped earlier and we get $T_{\mathrm{abs.\ phase}} < \tau_{\mathrm{res}}^{\mathrm{seg}}$. To minimize the bacterial growth, it is needed that $T_{\mathrm{abs.\ phase}} < \tau_{\mathrm{g}}$, which can also imply $T_{\mathrm{abs.\ phase}} < \tau_{\mathrm{abs}}$, provoking a loss of nutrients. When the phase duration is maximized to be $T_{\mathrm{abs.\ phase}} \simeq \tau_{\mathrm{g}}$, the nutrient loss is minimized.

- $T_{\mathrm{cleanphase}}$ the duration of the flushing phase. Since this phase is designed to reduce bacterial numbers, it should last as the residence time of bacteria in the tube $T_{\mathrm{clean\ phase}} \simeq \tau_{\mathrm{res}}^{\mathrm{per}}$. To be efficient in reducing bacterial numbers and avoid fast bacterial replication, this phase should last less than the bacterial growth time; therefore, fast flows should be preferred so that $T_{\mathrm{clean\ phase}} \simeq \tau_{\mathrm{res}}^{\mathrm{per}} \ll \tau_{\mathrm{g}}$.

##### C. Nutrient loss due to the bacterial threshold mechanism

How can we quantify the nutrient loss due to a too short absorption phase when the bacterial threshold mechanism is used? As shown in section IX, if nutrients are not leaving the tube due to advection, the nutrient concentration in the tube is given by $N_{\mathrm{in\ tube}} = N_0 \exp(-t/\tau_{\mathrm{abs}})$. This approximation is valid if we look at times when the nutrients mainly did not leave the tube, so for times $t < \tau_{\mathrm{res}}$. Both of our phases last at maximum as the respective residence times, so we can use this exponential decay to describe the system in good approximation. Therefore, the nutrients which do not get absorbed during the two phases and are lost are given by $N_{\mathrm{lost}} = N_0 \left(1 - \exp\left(-\left(T_{\mathrm{abs.\ phase}} + T_{\mathrm{clean\ phase}}\right)/\tau_{\mathrm{abs}}\right)\right)$ (assuming that bacteria consumption is always not relevant).

For the bacterial threshold case, we saw in the preceding section that the absorption phase should be $T_{\mathrm{abs.\ phase}} \simeq \tau_{\mathrm{g}}$, while the flushing phase is of the order of $\tau_{\mathrm{res}}^{\mathrm{per}}$. Substituting this, we obtain

$$N_{\mathrm{lost}} = N_0 \left(1 - \exp\left(-\frac{\tau_{\mathrm{g}} + \tau_{\mathrm{res}}^{\mathrm{per}}}{\tau_{\mathrm{abs}}}\right)\right) \tag{84}$$

Therefore, for the bacterial feedback case, the timescale ratio which governs the loss of nutrients is given by

$$\frac{\tau_{\mathrm{g}} + \tau_{\mathrm{res}}^{\mathrm{per}}}{\tau_{\mathrm{abs}}}. \tag{85}$$

The smaller this ratio, the bigger the loss and *viceversa*.

[1] J.D. Huizinga, S.P. Parsons, J.H. Chen, A. Pawelka, M. Pistilli, C. Li, Y. Yu, P. Ye, Q. Liu, M. Tong, *et al.*, Am. J. Physiol. Cell Physiol. **309(6)**, C403 (2015).
[2] S. Marbach, and K. Alim, Phys. Rev. Fluids **4**, 114202 (2019).
[3] M. Li, and J.G. Brasseur, J.Fluid Mech. **248**, 129 (1993).
[4] F. Hugenholtz, and W.M. de Vos, Cell. Mol. Life Sci. **75(1)**, 149 (2018).
[5] A. Pereira de Souza, R. Sieberg, H. Li, H.R. Cahill, D. Zhao, T.C. Araújo-Jorge, H.B. Tanowitz, and L.A. Jelicks, Parasitol. Res. **106(6)**, 1293 (2010).
[6] H. Ohkubo, T. Kessoku, A. Fuyuki, H. Iida, M. Inamori, T. Fujii, H. Kawamura, Y. Hata, N. Manabe, T. Chiba, *et. al*, Am. J. Gastroenterol. **108(7)**, 1130 (2013).
[7] T. Ishikawa, T. Sato, G. Mohit, Y. Imai, and T. Yamaguchi, J. Theor. Biol. **279**, 63 (2011).
[8] T.E. Moxon, O. Mihailova, O. Gouseti, P.J. Frye, and S. Bakalis, Engineering Digestion: Effect of Viscosity and Gastric Secretions on the Absorption of Nutrients, in Gums and Stabilisers for the Food Industry 18: Hydrocolloid Functionality for Affordable and Sustainable Global Food Solutions, (The Royal Society of Chemistry 2016), 264 .
[9] G. I. Taylor, Proc. R. Soc. Lond. A **219**, 186 (1953).
[10] G. I. Taylor, Proc. R. Soc. Lond. A **223**, 446 (1954).
[11] R. Aris, Proc. R. Soc. Lond. A **235**, 67 (1956).
[12] G. N. Mercer, and A. J. Roberts, SIAM J. Appl. Math. **50**, 1547 (1990).
[13] G.N. Mercer, and A.J. Roberts, Jpn. J. Ind. Appl. Math. **11**, 499 (1994).
[14] J. Gerritse, F. Schut, and J.C. Gotischal, Appl. Environ. Microbiol. **58(5)**, 1466 (1992).
[15] K.A. Johnson, and R.S. Goody, Biochemistry **50(39)**, 8264 (2011).
[16] G. Canbolat, H.A. Kose, A.l. Yildizeli ,and S. Cadirci, Analytical and numerical solutions of the 1D advection-diffusion equation in 5th International conference on advances in mechanical engineering (Istanbul 2019).
[17] P.W.M. Janssen, R.G. Lentle, P. Asvarujanon, P. Chambers, K.J. Stafford, and Y. Hemar, J. Physiol. **582(3)**, 1239 (2007).
[18] K.E.Barrett, S.M. Barman, J. Yuan, and H.L. Brooks, Ganong's Review of Medical Physiology (McGraw Hill Education, New York, 2016), Twenty-fifth edition.
[19] J.D. Huizinga, J.H. Chen, Y. F. Zhu, A. Pawelka, R.J. McGinn, B.L. Bardakjian, S.P. Parsons, WA. Kunze, R.Y. Wu, P. Bercik, *et al.*, Nat. Commun. **5**:3326 (2014).
[20] K.E.Barrett, S.M. Barman, J. Yuan, and H.L. Brooks, Ganong's Review of Medical Physiology (McGraw Hill Education, New York, 2016), Twenty-fifth edition.
[21] B. Waclawiková, A. Codutti, K. Alim, and S. El Aidy, Gut Microbes **14(1)**, 1997296 (2022).
